# Local Dendritic Landscape of Mouse V1

**DOI:** 10.1101/2025.07.10.664234

**Authors:** Nelson D. Medina, Anastasia Sorokina, Narayanan Kasthuri

## Abstract

The principles that govern where excitatory synapses form on cortical pyramidal neurons remain unclear. A long-standing hypothesis is that connectivity mirrors neuronal geometry—with apical dendrites dominating in layer 1 and basal dendrites in deeper layers. Leveraging the MiCRONS cubic-millimetre serial electron-microscopy dataset ([1, 2]), we mapped 4968 synapses onto reconstructed apical and basal arbors across layers 1–6 of mouse visual cortex. Contrary to a simple gradient model, synaptic targeting is overwhelmingly biased toward basal dendrites in layers 2/3–6, even where apical shafts are abundant. Layer 1 is the sole exception, exhibiting the expected apical bias. Basal predominance scales with local soma density and the relative arbor length available for contact, such that 66% (61101/92445) of all synapses land within 130 µm of the soma, placing most excitatory drive near the cell body. Axonal analysis revealed specificity beyond mere geometric opportunity: individual axon segments preferentially innervated either basal or apical compartments far more often than predicted by chance, indicating compartment-selective wiring rules. Together, our results show that cortical connectivity is shaped by both neuronal geometry and axon-level targeting preferences, redefining how pyramidal cells integrate information across layers and apical and basal dendrites.

## 1 Introduction

Understanding how synaptic wiring shapes neuronal responses is a central problem in neuroscience. In the cerebral cortex, pyramidal neurons—the principal excitatory cells—integrate inputs in two morphologically and functionally distinct dendritic domains: apical and basal [3]. Each domain receives characteristic input streams and supports different computations [4, 5, 6, 7, 8, 9].

Apical dendrites extend toward the pial surface, where they are contacted by long-range thalamic afferents [10, 11] and feedback projections from higher cortical areas, integrating contextual information and top-down signals [12, 13, 9]

Basal dendrites, which ramify near the soma, mainly receive local cortico-cortical input that shapes a neuron’s receptive field and feed-forward sensory drive [14, 15, 16, 17].

Although these functional roles are well established, the quantitative rules that govern apical versus basal innervation across cortical layers remain unclear. A simple geometric hypothesis predicts a smooth layer-by-layer gradient—apical targeting in layer 1, basal targeting in layer 6. But do real circuits follow this monotonic trend, or do they exhibit more intricate, compartment-specific patterns? Answering this requires synapse-level data spanning multiple layers—data that have only recently become available.

The advent of large-scale serial electron microscopy (EM) datasets has revolutionized our ability to study cortical circuitry [18]. In this study, we used the MiCRONS *mm*^3^ dataset to investigate how apical and basal dendrites are innervated across layers of primary visual cortex (V1) [1, 19]. Specifically, we asked how patterns of synaptic input relate to both the cortical layer and the geometry of individual pyramidal neurons.

We found that instead of innervation following a monotonic gradient, with apical targeting dominating in superficial layers (e.g., layer 1) and basal targeting increasing in deeper layers, our analysis revealed a more complex picture:

1. We found non-monotonic patterns of dendritic innervation that could not be explained solely by the relative lengths of apical and basal dendrites in each layer.
2. We observed that postsynaptic targeting by axons was not reliably predicted by the local availability of dendritic branches. Instead, the distribution of apical versus basal inputs appeared to depend on a combination of soma density and neuronal geometry.
3. Finally, we found that most synapses onto pyramidal neurons are located proximally to the soma, suggesting a spatial bias that may reflect functional constraints on input integration.

Together, these results show that cortical connectivity is governed by both geometric constraints and compartment-specific targeting preferences, challenging the traditional gradient model and refining our understanding of how layered architecture supports distributed computation.

## 2 Results

We first investigated the distribution of basal and apical synapses in Layer 4 (5-micron radius, Fig. 1D). We visually identified synapses using vesicle clouds in the presynaptic terminal and densities on the postsynaptic terminal (PSD) (Fig. 1D, highlighted spines in the right panels) and traced the postsynaptic path to the cell body (Fig. 1E, automatic segmentation from MiCRONS)[20, 2]. This resulted in a sample of 35 synapses. Two synapses belonged to nonpyramidal neurons (putatively inhibitory), 5 were untraceable back to their somas, and 28 were spine synapses from pyramidal neurons (Fig. 1F). We found that most synapses were on nearby basal dendrites (24/28). Only 3 synapses were formed on neurons whose cell bodies outside layer 4 (Fig. 1F), which belonged to layer 5 and layer 6 neurons.

**Figure 1:**
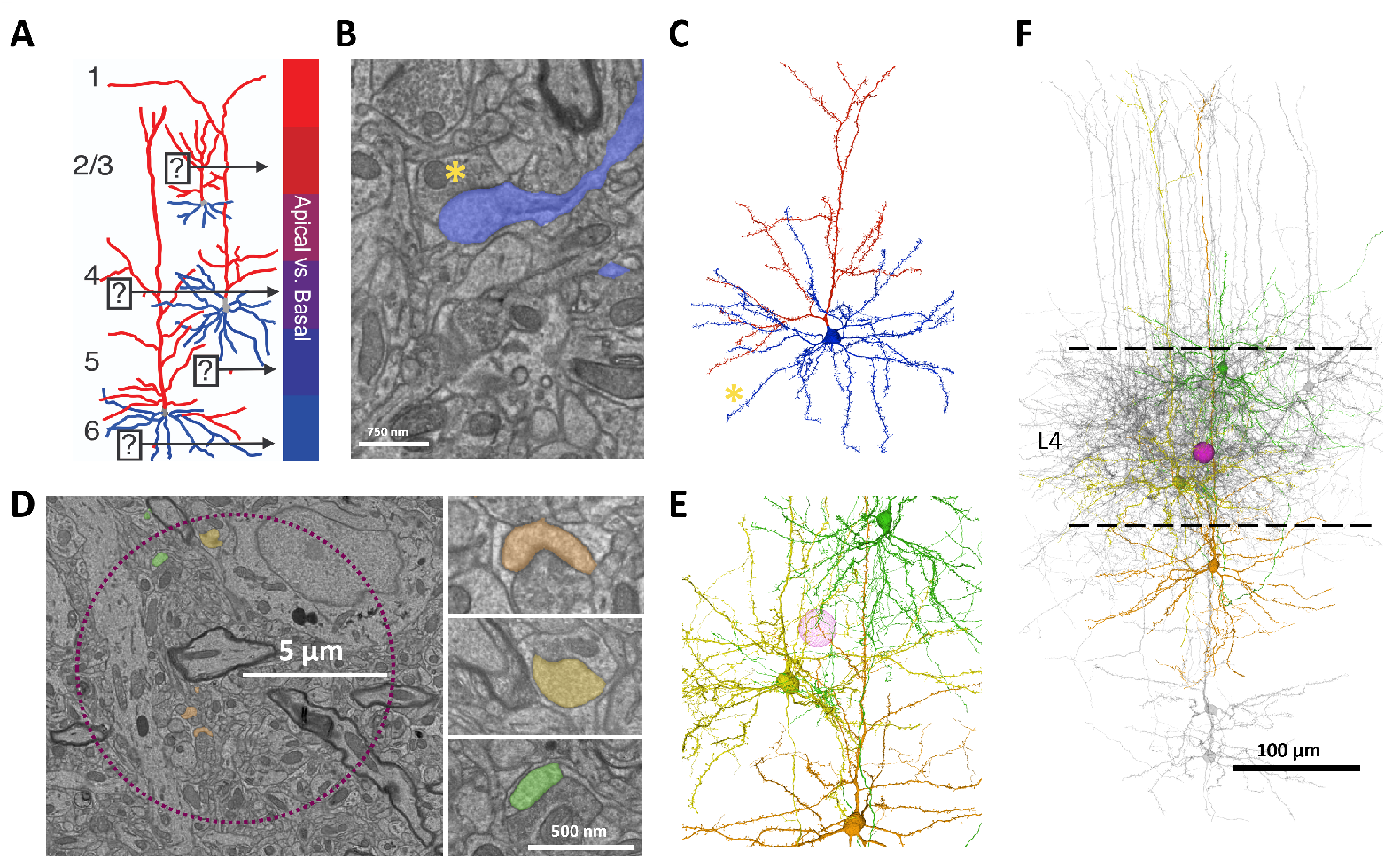
Apical and Basal categorization of synapses. **A:** Schematic representation of apical and basal dendrites across cortical layers. Apical dendrites are shown in red and basal dendrites in blue. **B:** Electron microscopy image of synapse. An example synapse is shown with a yellow asterisk over the presynaptic terminal, and postsynaptic terminal and dendrite highlighted in blue. **C:** A 3D reconstruction of the neuron and synapse in B (marked by a yellow asterisk). Basal dendrites are colored blue, apical dendrite colored red. **D:** Manual sampling of synapses in a 5-Micron radius. Three example synapses sampled from within the search region (purple circle). **E:** 3D reconstructions of the three neurons from synapses shown in D. **F:** Reconstructions of 28 neurons whose synapses were sampled from within the area shown in D.

Next, we asked whether the proportion of apical to basal synapses varied by layer. A combination of strategies was used to sample across all layers of the V1 mm^3^ dataset. First, three locations (yellow triangle, circle, and square, for layers 1, 2/3, and 5, respectively, Fig. 2) were sampled within the same limited region as before (Fig. 1D). The second approach used randomly generated coordinates along a thin (5 by 5 micron) column that spanned all cortical layers. The coordinates were then used to constrain a manual search for synapses. In sum, the combined approach yielded 846 synapses, of which 16.2% were untraceable back to their soma (26/137) or belonged to non-pyramidal neurons (111/137). Of the remaining 689 synapses, we found that 63% of the synapses occur in the basal dendrites across all cortical areas (435/689). We then binned the data into 10 bins by cortical depth (Fig. 2 A). The median percent basal for synapses in this binning was 73.5% basal. At most depths, the proportion of basal synapses deviated above the expected 63% average, with two notable exceptions: layer 1 and the intersection of layer 4 and layer 5. In layer 1, 93% of the synapses occur in the apical dendrites (yellow triangle, Fig. 2A). Between layers 4 and 5, the percentage of basal synapses sampled was 53%. For the vast majority of cortical space (excluding these two exceptions), basal dendrites dominated synaptic availability (74.5 % basal synapses). For some layers, this percentage deviated significantly (75-80% basal) from the simple average of 63% (Fig. 2A).

**Figure 2:**
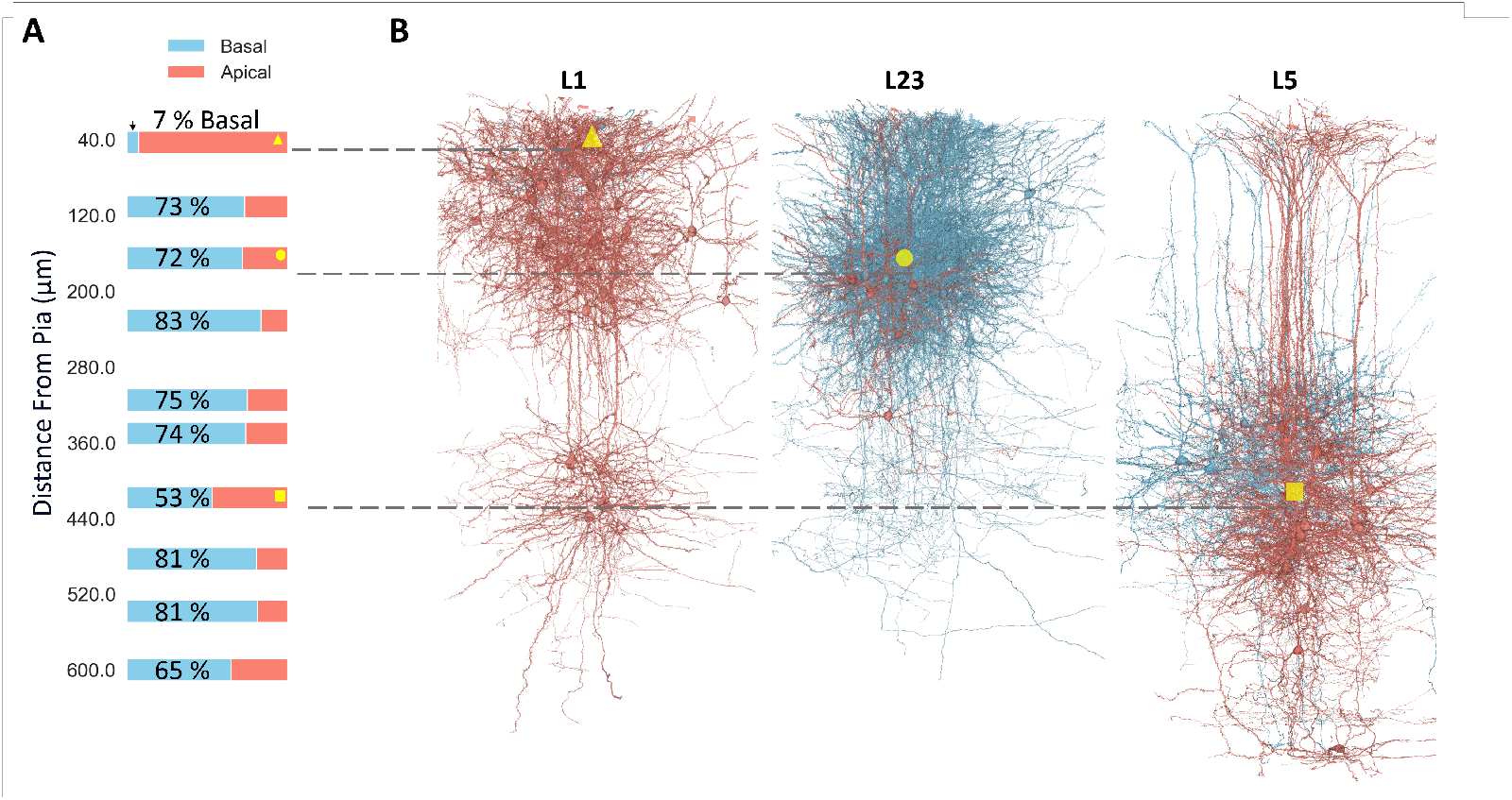
Basal synapses dominate across cortical layers except in layer 1 and the L4-L5 border. (A) Percentage of synapses occurring on basal (blue) versus apical (red) dendrites across cortical depth, binned in 10 intervals from pia to white matter. In most bins, the proportion of basal synapses exceeded the overall average of 63%, with a median of 73.5%. Notable exceptions include layer 1 (yellow triangle), where 93% of synapses were apical, and the L4-L5 border (yellow square), where basal synapses comprised only 53%. (B) Segmented pyramidal neurons surrounding three sampled coordinates (yellow triangle, circle, and square for L1, L2/3, and L5, respectively) are shown. Neurons are colored according to the dendritic compartment receiving a synapse near the sampled coordinate: red for apical and blue for basal. Only the local synapse near the sampling point determines the neuron’s color, not its overall dendritic morphology.

We also find that most synapses are made on nearby neurons, often within the same layer as the synapse (layer 4 synapses in Fig. 1F and layer 2/3 in Fig. 2B, for example). This was expected for basal dendrites, given that their length is only a few hundred microns [21, 8, 9, 3, 10, 18, 1]. However, this was also true for apical synapses. For example, in layer 1 where most apical dendrites terminate and arborize, we found that most of the synapses on apical dendrites came from nearby neurons in layer 2/3. In our sampling, only 7% of synapses came from outside layer 2/3 (4/54), and all four outside synapses came from the apical branches of layer 5 neurons.

This prompted us to ask how far synapses were made on average from neuronal soma. We measured the path length from a synapse back to the soma for every detected synapse (see Methods) on a neuron (Supplementary Fig. 1A). We did this algorithmically for 5 neurons from 5 classes (L6, L5b, L5a, L4, and L2/3) for a total of 25 neurons (100 synapse paths shown for one of each neuron class, Supplementary Fig. 1B). We found that most synapses are 100–150 microns from the soma, by cable length (average = 130 microns, median = 105 microns, standard deviation = 103 microns) (Supplementary Fig. 1C). This was true for all neuron types, including those with vast apical arborizations. Layer 5 and layer 4 neurons, for example, contain secondary peaks in length distributions corresponding to distal apical branches. We then did this again for another 5 neurons from each layer (L2/3, L4, L5 and L6, Supplementary Fig. 2) to determine the distributions of path lengths from synapse to soma of apical synapses. We found that the average synapse cable length for apical synapses were: 245.2 (Std 15.09) for L5, 123.2 (Std 4.40) for L4, 121.4 (Std 4.63) for L2/3, and 183.4 (Std 31.93) for L6. This is consistent with the average total synaptic cable lengths we find, suggesting most dendrites in cortex are more local than previously appreciated (Supplementary Fig. 2)

Next, we extended our manual sampling with algorithmically determined apical-basal synapse identification. We used the automatically identified synapse table in the MiCRONS dataset [2] to select synapses within a 5 by 5 micron column that spanned all cortical layers (4,122 synapses, Fig.3A). We then skeletonized the 3D reconstructions of neurons whose synapses were selected (neurons without a nucleus were excluded). For each neuron, we broke the skeleton into its main component branches and assigned synapses onto their nearest branch. Finally, we identified the apical dendrite by finding the synapse center of mass for each branch and choosing the one closest to the pia surface (Fig. 3B, top panel). With this methodology, we found that 63% of the synapses on an average pyramidal neuron are on basal dendrites (Fig. 3B, bottom panel). This is consistent with previous observations of apical and basal dendritic lengths [10]. As before, we found that most synapses are relatively local for both basal and apical synapses. Plotting the location of the soma against the location of the synapse for the basal and apical synapses reveals very similar linear distributions (Fig. 3C), reflecting the proximity of most synapses for a given neuron. A secondary distribution can be seen for apical synapses, which represents a small number of synapses on the main apical branch, the collateral branches, and the arborization that occurs near layer 1. The proportions of apical and basal synapses from this approach agreed with our manual sampling and identified the same two exceptions to basal synapse dominance (layer 1 and threshold of layers 4 and 5, Fig. 3D).

**Figure 3:**
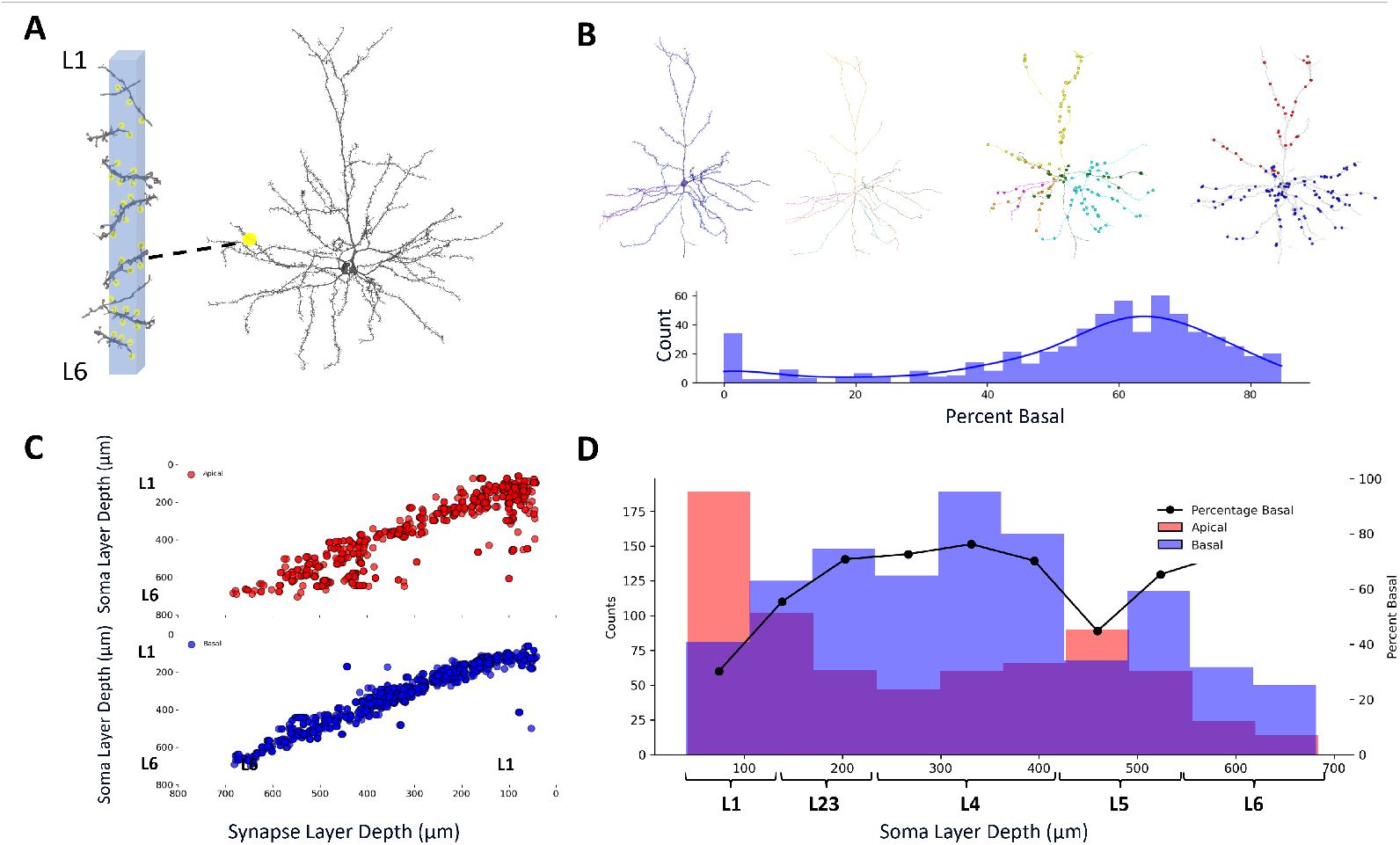
Automated classification of apical and basal synapses confirms basal dominance and local connectivity. (A) Synapses within a 5 *µ*m *×* 5 *µ*m column spanning all cortical layers were extracted from the MiCRONS dataset (yellow dots), and neurons receiving those synapses were reconstructed and skeletonized. (B) *Top*: Example skeletons of neurons with distinct primary branches used to identify the apical dendrite based on proximity to the pia. Synapses were assigned to their nearest branch and the apical branch was defined as the one with a center of mass closest to the pia. *Bottom*: Distribution of the percentage of basal synapses across neurons, with a mean of 63% basal, consistent with previous analyses and dendritic length measurements. (C) Soma depth plotted against synapse depth for all apical (red) and basal (blue) synapses shows that most synapses occur locally, near the soma. A secondary distribution of apical synapses reflects inputs onto apical tufts and collaterals near layer 1. (D) Histogram showing apical (red) and basal (blue) synapse counts as a function of soma depth. A black line indicates the percent of synapses that are basal in each depth bin. As in manual sampling, layer 1 and the border between layers 4 and 5 show deviations from the otherwise consistent basal dominance across cortical layers.

Given the relative proximity of synapses to their respective neuron’s soma, we next examined how neuron density relates to the percentage of basal synapses across cortical layers. Neuron density varies substantially between layers (Fig. 4A, distribution by class and examples) and across species [22, 23]. In mouse V1, pyramidal neuron density follows a bimodal distribution, with peaks in layer 4 and layer 6, and a clear dip in layer 5 (Fig. 4) [24, 25, 26]. In contrast, inhibitory neurons show an inverse pattern, peaking in layer 4 (Fig. 4). Notably, we find that the distribution of excitatory cell bodies (gray density, Fig. 4B) closely tracks the proportion of basal synapses across layers (blue line, Fig. 4B).

**Figure 4:**
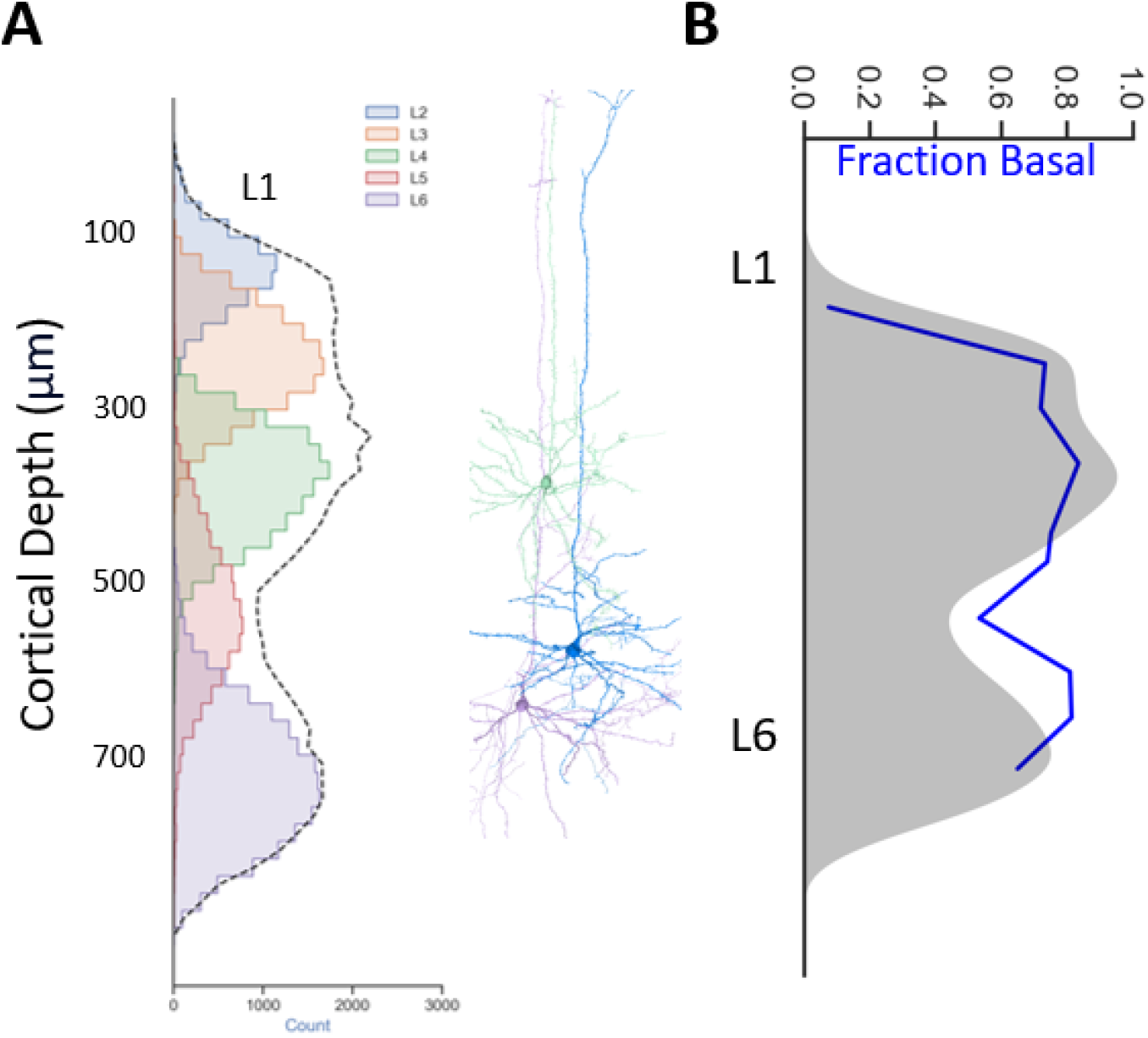
Pyramidal neuron density across layers tracks the distribution of basal synapses. (A) Left: Histogram of pyramidal neuron soma counts as a function of cortical depth in mouse V1. Layer boundaries are overlaid and color-coded. The dashed black line shows the overall density of excitatory neurons, which peaks in layers 4 and 6 and dips in layer 5. Right: Example pyramidal neuron reconstructions from different layers highlight diversity in morphology and soma depth. (B) The fraction of synapses assigned to basal dendrites (blue line) closely follows the distribution of excitatory neuron density (gray shaded area), suggesting that local neuron density helps shape the relative abundance of basal synaptic input across cortical depth.

These observations suggest that synaptic distributions may largely reflect local neuron density. However, local availability alone may not fully explain synaptic targeting—there could be additional presynaptic mechanisms that guide connectivity in a more selective or structured way.

To test whether axons in V1 follow or deviate from Peter’s rule, we used two complementary approaches. First, we manually traced individual pyramidal neuron axons and identified the postsynaptic targets of their first few boutons (up to 10 synapses; Supplementary Fig. 4A). Second, we selected random axon segments not traced back to their somas (Fig. 5A) and manually validated as many synapses as possible, ensuring continuity of the axon and agreement between human annotators (10–20 synapses; Fig. 5A).

**Figure 5:**
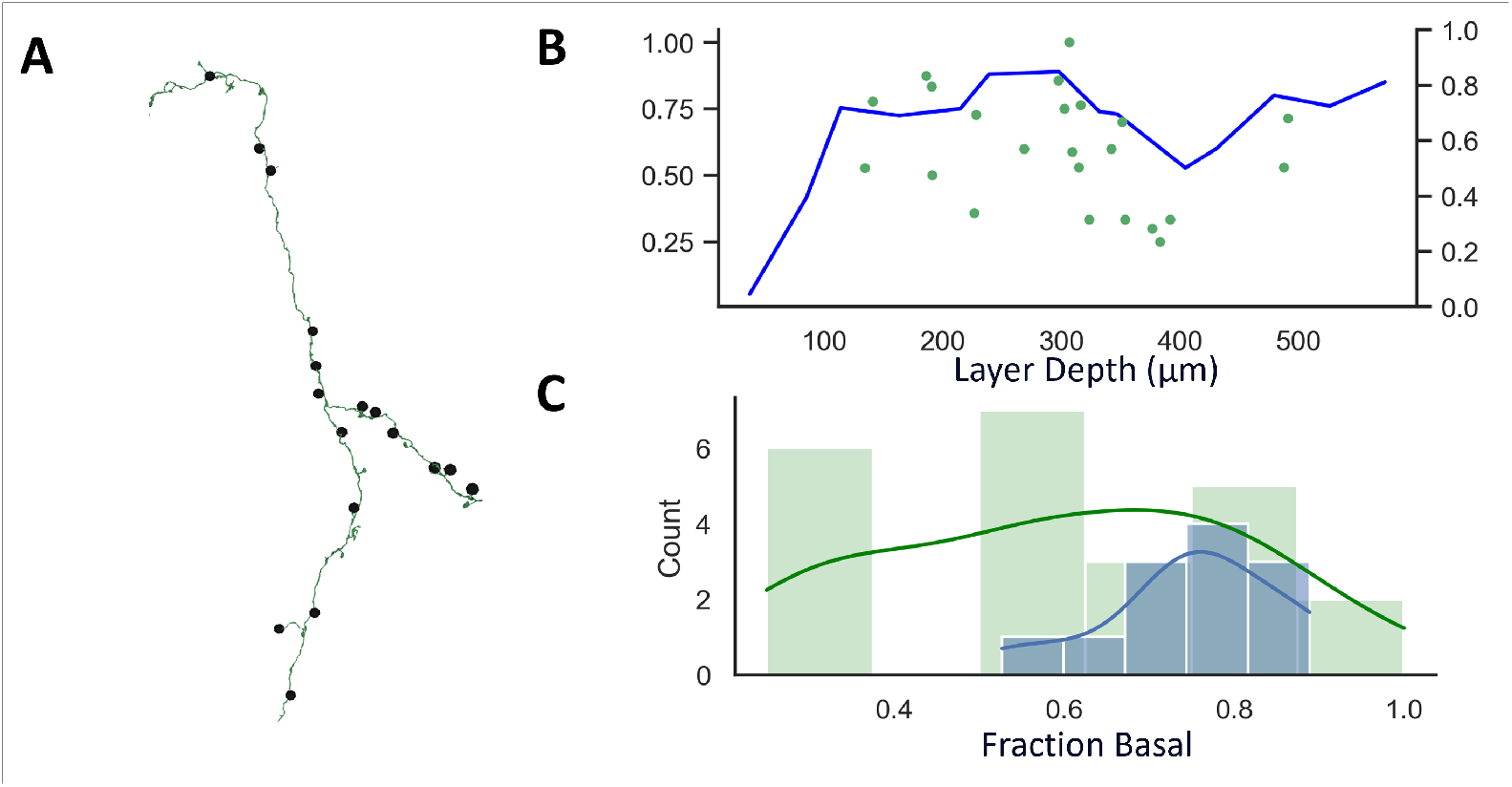
Detached axon segments reveal variability in postsynaptic targeting and deviations from population-level basal bias. (A) Example of a manually validated axon segment (green) with synapse locations indicated (black dots). Only segments with continuous axonal morphology and high annotator agreement were included. (B) Fraction of synapses targeting basal dendrites (blue line, right axis) plotted by cortical depth for each axon segment (green dots, left axis), showing overall basal preference with substantial variability. (C) Histogram of the fraction of basal synapses for all 23 axon segments (blue bars), compared to the expected distribution of basal synapses by cortical depth (green bars). While most axons show a strong basal bias, a subset (7/23) fall below 50% basal, indicating that axonal targeting does not uniformly follow layer-level synapse distributions. A KS test confirms a significant difference between axonal and population-level distributions (*p* = 0.035).

In the first approach, we analyzed 20 neurons with a total of 70 synapses (average 3.5 synapses per axon, standard deviation 2.9). Consistent with previous findings in entorhinal cortex (path-length-dependent synapse sorting) [27], we observed that proximal axon boutons formed synapses with interneurons about half the time (31/70 synapses on nonpyramidal neurons, data not shown). Among the remaining synapses, the majority targeted basal dendrites (22 basal synapses), followed by apical dendrites (10 synapses), and somata (3 synapses). All synapse annotations were cross-validated by at least two human experts. We then calculated presynaptic and postsynaptic path lengths back to their respective somas (same method as in Supplementary Fig. 1A) and found that the Euclidean distance between the somas of connected neurons (Supplementary Fig. 4B) significantly correlated with both presynaptic (p value = 0.0001, Pearson’s correlation) and postsynaptic (p value ¡ 0.0001, Pearson’s correlation) cable lengths (Supplementary Fig. 4C, D). Interestingly, pre- and postsynaptic cable lengths did not correlate with each other (p value = 0.11, Pearson’s correlation, data not shown).

For the second approach, we analyzed 23 ‘detached’ axon segments—those not traced back to a soma—capturing a total of 283 synapses (mean 12.3 synapses per segment, standard deviation 4.98). Most segments showed a clear bias toward targeting basal dendrites (blue bars, Fig. 5D). However, this bias was not universal: 7 out of 23 axon segments had a basal preference below 50% (green data points, Fig. 5C), deviating substantially from both the average neuron-level basal bias (63%) and the average layer-level bias (75%). When comparing these axonal distributions to the baseline basal synapse percentages by layer (excluding L1), we found a significant difference (p value = 0.035, estimated power = 0.67, KS test, Fig. 5C).

Lastly, to extend our analysis beyond the limited manually traced axon boutons—and to address the labor-intensive nature of manual tracing—we applied an algorithmic approach (similar to the method used in Fig. 2). Specifically, we identified, at most, the first 20 boutons along each axon based on path length, and measured both the distance between connected neuron somas and the postsynaptic dendritic path lengths (Fig. 6B). This analysis was restricted to neurons whose 3D reconstructions contained substantial portions of their local axon. We selected 30 such neurons (INFO ON NEURONS) and algorithmically identified a total of 606 synapses.

**Figure 6:**
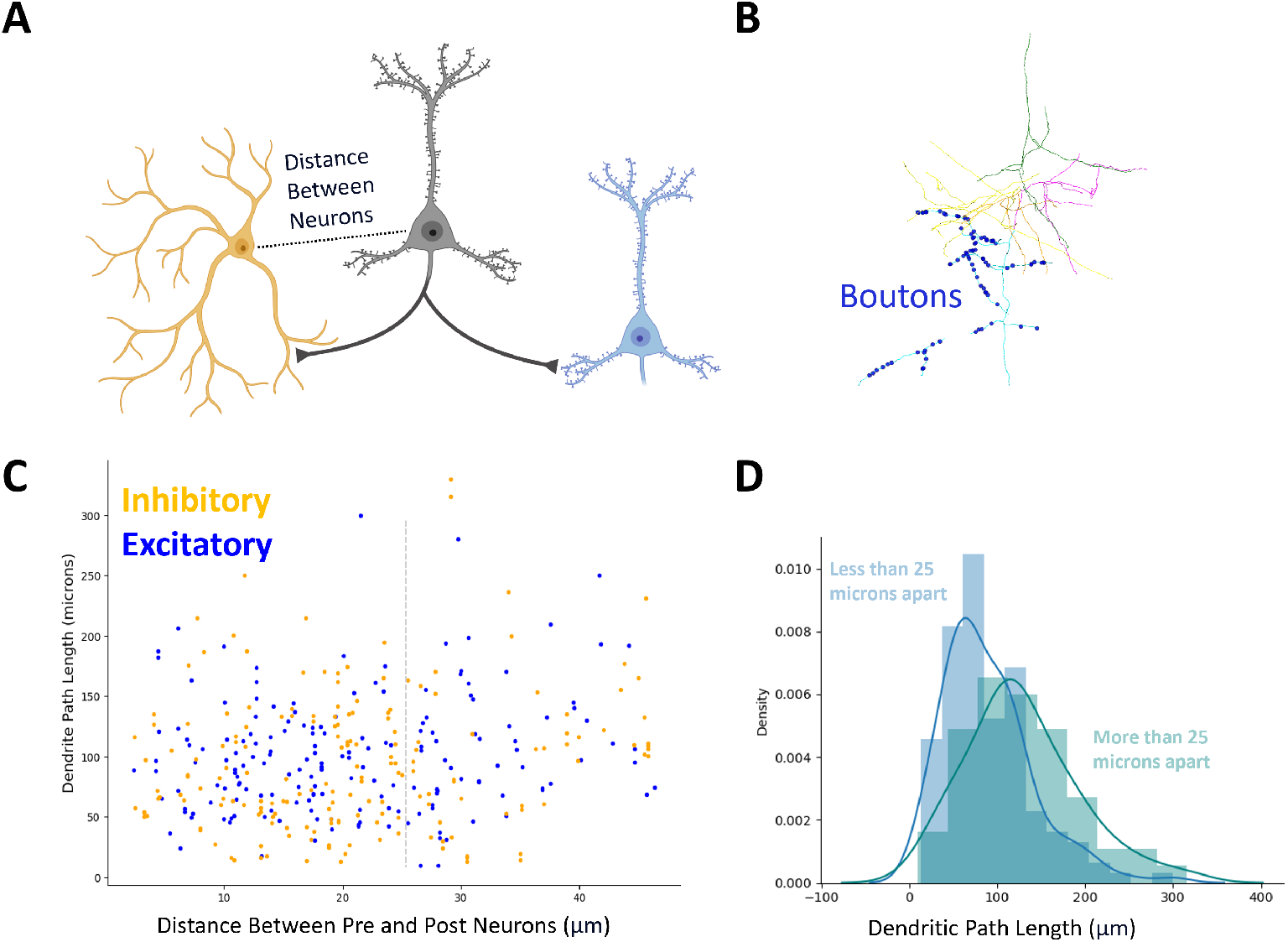
Automated bouton mapping reveals proximity bias in inhibitory connectivity and postsynaptic path length. (A) Schematic illustration of the analysis: for each neuron, up to 20 boutons were mapped and both intersomatic distances and postsynaptic dendritic path lengths were measured. (B) Example axon reconstruction with boutons (dark blue dots) identified algorithmically along its length. (C) Scatter plot showing dendritic path lengths as a function of distance between presynaptic and postsynaptic neuron somas, color-coded by postsynaptic cell type (orange: inhibitory, blue: excitatory). Inhibitory connections tend to occur more proximally, with 56% of inhibitory synapses within the first 10 boutons. (D) Histogram of dendritic path lengths grouped by intersomatic distance: pairs less than 25 *µ*m apart (blue) have significantly shorter postsynaptic path lengths than those farther apart (green). Mean dendritic path lengths were 89.3 *µ*m and 128.1 *µ*m for close and distant pairs, respectively.

Using the MiCRONS neuron classification tables [20, 2], we were able to categorize 76% of the connected partners as excitatory or inhibitory. This dataset once again reveals a strong preference for proximal inhibitory connections: 56% of inhibitory synapses occurred within the first 10 boutons, and 43.1% within the second 10 boutons (Fig. 6C).

We also observed a relationship between intersomatic distance and postsynaptic path length. Neuron pairs whose somas were closer than 25 microns apart had significantly shorter postsynaptic dendritic path lengths compared to those further apart (Fig. 6C, D). Specifically, close neuron pairs had an average dendritic path length of 89.3 microns (Fig. 6D, blue histogram), whereas more distant pairs averaged 128.1 microns (Fig. 6D, green histogram). Notably, we did not observe any significant differences in postsynaptic path lengths between excitatory and inhibitory target neurons (Supplementary Fig. 5).

## 3 Discussion

The role of neuronal morphology and dendritic compartmentalization in circuit function has been extensively studied and is widely recognized as a foundational aspect of neuronal computation [21, 28, 3, 5, 29, 30, 31]. Many small, self-contained circuits have been well-characterized, with structural [18, 1, 20, 2, 10] and functional [14, 15, 16, 17] properties that map onto behavior, sensation, and perception [12, 13]. However, traditional circuit diagrams often omit the broader structural context in which these microcircuits are embedded. This motivated us to investigate the ‘background’ architecture in which specific connections are situated. Given the well-established roles of dendritic compartments [21, 28, 3, 5, 29] and cortical layer organization, we specifically asked how synaptic inputs onto basal and apical dendrites are spatially distributed across the cortex.

We found that basal dendrites—and their associated synapses—constitute 63% of the average pyramidal neuron in V1, aligning with previous reports [4, 15, 16]. When considering all pyramidal synapses within a 5 by 5 micron voxel, the visual cortex as a whole also comprises approximately 63% basal synapses. However, the laminar distribution of apical and basal synapses was far from uniform. As expected, layer 1—where apical dendrites arborize and receive input from other cortical areas—is dominated by apical synapses (93% apical). At the border of layers 4 and 5, where oblique branches from layer 5 neurons emerge, the apical-to-basal ratio approaches parity (50% apical). Outside of these two zones, basal synapses dominate; notably, layer 4 shows a strong basal bias, with roughly 80% of synapses targeting basal dendrites. These findings suggest that apical synapses in most cortical regions are in the minority, potentially posing challenges to the isolation and maintenance of distinct computational compartments. Yet, we observed that individual axon segments often diverged markedly from the expected background distributions—for example, some axons almost exclusively targeted apical dendrites within layer 4 (Fig. 5F).

These results are consistent with previous experimental findings [10, 11] and computational models describing differential thalamo-cortical innervation in primary somatosensory cortex. While those studies did not explicitly frame the pattern as a basal bias, the majority of thalamo-cortical axons were shown to preferentially target basal dendrites. Whether this reflects simple dendritic availability in layers 4 and upper 5, or instead the action of specific axonal targeting rules, remains unresolved. Importantly, in those previous studies, thalamo-cortical axons did not deviate from the expected background targeting rates—unlike the axonal segments we analyzed, which frequently exhibited significant biases. These observations underscore the need for future experiments to disentangle synaptic specificity from constraints imposed by dendritic availability.

Our dataset also represents a stable snapshot of the mature cortex—an adult state shaped by both genetic programming and sensory experience. In early development, apical dendrites make up the majority of pyramidal neuron dendritic length in mice [32]. This raises the possibility that the apical-to-basal balance may shift toward basal dominance over time as a consequence of experience. Supporting this idea, sensory deprivation in young mice has been shown to selectively impair basal dendritic development while leaving apical dendrites relatively unaffected [8]. This would result in a different landscape of dendritic availability and, consequently, different cortical processing dynamics.

One of the most unexpected findings was the prevalence of synapses located proximally to the soma (within 100–120 microns). While this is partly explained by the branching patterns of basal and proximal apical dendrites, we had anticipated a stronger contribution from apical tufts in layer 1. Instead, this effect appears to be counterbalanced by increasing neuron density in deeper layers (Fig. 3D and Fig. 4D). Furthermore, we found that both presynaptic and postsynaptic cable distances for the earliest axonal boutons were significantly correlated with the physical distance between neuron somas. This suggests that neurons in close spatial proximity tend to form structurally and electrically advantageous connections. Such proximity-based connectivity provides a plausible mechanistic basis for previous functional studies showing that nearby neurons often have highly correlated activity patterns [14], a phenomenon also evident in the MiCRONS dataset (Supplementary Fig. 6). An alternative explanation could be that these neurons receive common input. Of course, these two hypotheses are not mutually exclusive and may reflect a feedback mechanism in which proximity facilitates shared inputs, and vice versa.

## 4 Conclusion

In summary, our analysis of synaptic connectivity in mouse V1 reveals that the organization of apical and basal dendritic inputs is more complex than a simple layer-by-layer gradient. While the overall synapse distribution reflects the dominant role of basal dendrites in cortical processing, local deviations—driven by neuronal geometry, soma density, and axon-specific targeting—highlight a nuanced landscape of connectivity. These findings challenge the assumption that local dendritic availability solely dictates synaptic placement, and instead point to additional rules or biases shaping synaptic networks. The prominence of proximal synapses and the spatial dependence of connection patterns further suggest that cortical wiring supports efficient, compartmentalized computation that balances structural constraints with functional specificity. Moving forward, integrating developmental, molecular, and functional datasets with large-scale ultrastructural reconstructions will be critical for uncovering the mechanisms that govern these wiring patterns and their implications for perception and behavior.

## 5 Methods

### Dataset and Data Acquisition

#### Dataset: Mouse Visual Cortex Dataset

Data was accessed via the CAVE (Connectome Annotation Versioning Engine) [2] system and CloudVolume, which provide tools for efficient data processing and analysis. This dataset was made publicly available by the MICrONS Consortium via Microns Explorer, as part of an effort to map neural circuits. Manual synapse identification and dendritic categorization was performed by, and based on agreement of, two human experts. Segmentation versions v343 and v983 were used in our analysis.

#### Data acquisition protocols and preprocessing

Data acquisition for the MiCRONS dataset involved serial sectioning of cortical tissue, followed by imaging with a multi-beam scanning electron microscope. The resulting images were aligned and stitched to create a continuous volume, which was then segmented [2] to identify individual neurons, synapses, and other cellular components. Automated and manual annotation processes refined these segmentations to ensure accuracy and completeness (this is still ongoing).

### Synapse Identification and Classification

Synapse identification in the MiCRONS (mouse V1) dataset involves both automated detection algorithms and manual verification by expert annotators. Synapses are identified based on the presence of preand postsynaptic densities, vesicle clouds, and other ultrastructural markers observable in EM images. These classifications are included in tables accessible from CAVE.

#### Criteria for classifying synapses as basal, apical, or non-pyramidal

Synapse categorization was achieved using both automated tracing algorithms and manual tracing. The algorithmic solution involved skeletonizing the 3D meshes and then separating the skeletons into their main component branches (using the soma coordinate as a central point) and then assigning synapses to the nearest skeleton branch. Branches with fewer than 80 synapses were deemed as likely errors and excluded (axons were excluded this way). Then the synapse center of mass was identified for all branches, and the branch with center of mass closest to the pia surface was classified as the apical branch. All remaining branches were classified as basal. As before, any secondary branches attached to a main branch were classified as the main branch (e.g. synapses on oblique apical branches were classified as apical).

Two experts (co-first authors) manually identified and classified 854 synapses. Synapses were classified based on their location on the dendritic tree. Basal synapses were those found on primary dendrites stemming from the soma which did not extend towards the pia surface (non-apical), and apical synapses were those found on the main vertical dendrites of pyramidal neurons. Non-pyramidal synapses were identified on dendrites belonging to neurons that neither expert classified as having typical pyramidal morphology. Any synapse on either dendritic category, whether on the shaft or collateral branches, was included. All synapses that could not be traced back to their soma due to leaving the volume, or imaging artifacts, were excluded and marked as errors.

For analysis of individual axon segments, disconnected axon segments were identified via the neuroglancer MiCRONS explorer and manually traced to identify boutons. Three human experts validated each manual tracing and classified the postsynaptic dendrite as basal or apical.

#### Dendritic Path Length Analysis

Synapse-to-soma tracing involved following the postsynaptic dendrite from identified synapses back to their cell bodies. An algorithmic solution used the coordinates of identified synapses to find the closest point on the 3D reconstructed mesh and found the shortest path within the mesh from the synapse to the soma coordinates. All meshes used were down-sampled to 10% of their original size (90% decimation) for computational ease. Soma and synapse coordinates were obtained from opensource tables accessible via CAVE. This process was done for every synapse on 25 neurons (5 from 5 distinct layers).

For the analyses of the first few synapses on an axon human experts manually traced axons starting at the soma and identified the first 1-10 synapses on 28 neurons. Then, for each synapse, the postsynaptic side was manually traced back to the postsynaptic neuron’s soma and classified as basal or apical. Finally, the path length was calculated for the presynaptic and postsynaptic neurons (described above).

### Quantitative Analysis of Synaptic Distribution

#### Methods for calculating the proportion of basal and apical synapses

The per neuron proportion of basal and apical synapses was calculated by counting synapses on each type of dendrite and normalizing these counts by the total number of synapses. This analysis is performed across different cortical layers to identify patterns in synaptic distribution.

#### Binning data by cortical depth and statistical analysis

Synapse category was binned by cortical depth to examine synaptic distributions within specific cortical layers. Statistical analyses, including histograms and density plots, are used to visualize and quantify the distribution patterns. Statistical comparisons of distributions used the Kolmogorov-Smirnov (KS) test unless otherwise stated.

---

**Figure S1:**
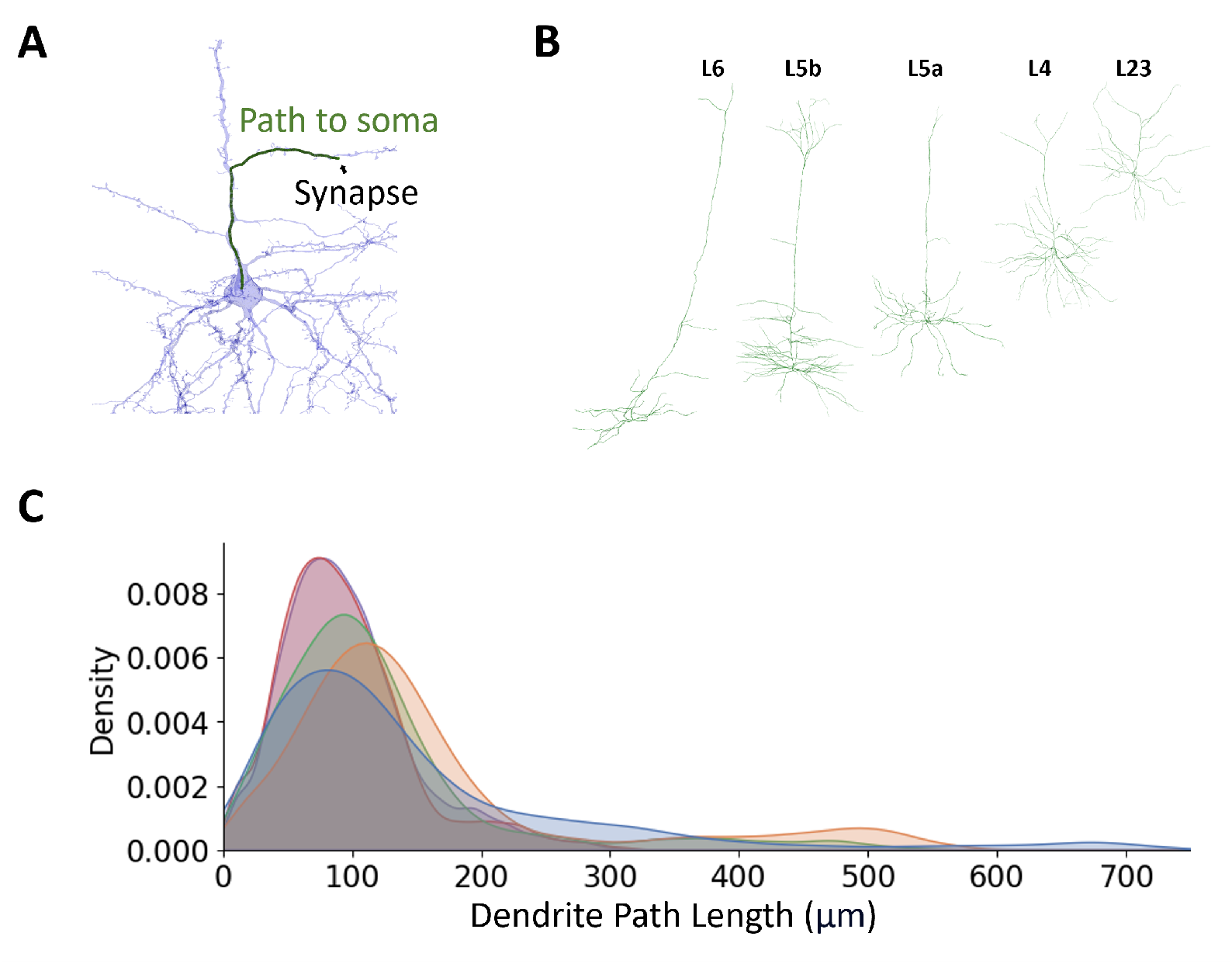
Most synapses are located within 100–150 *µ*m of the soma by dendritic path length. (A) Example neuron with a synapse (black dot) and the cable path traced back to the soma (green line) to compute dendritic path length. (B) Example reconstructions of 5 representative neurons (1 from each class: L6, L5b, L5a, L4, and L2/3), each with 100 synapse paths analyzed. (C) Density plots showing dendritic path length distributions across all neurons by class. Most synapses occur 100–150 *µ*m from the soma. Neurons with large apical trees (e.g., L5 and L4) show secondary peaks corresponding to distal apical branches.

**Figure S2:**
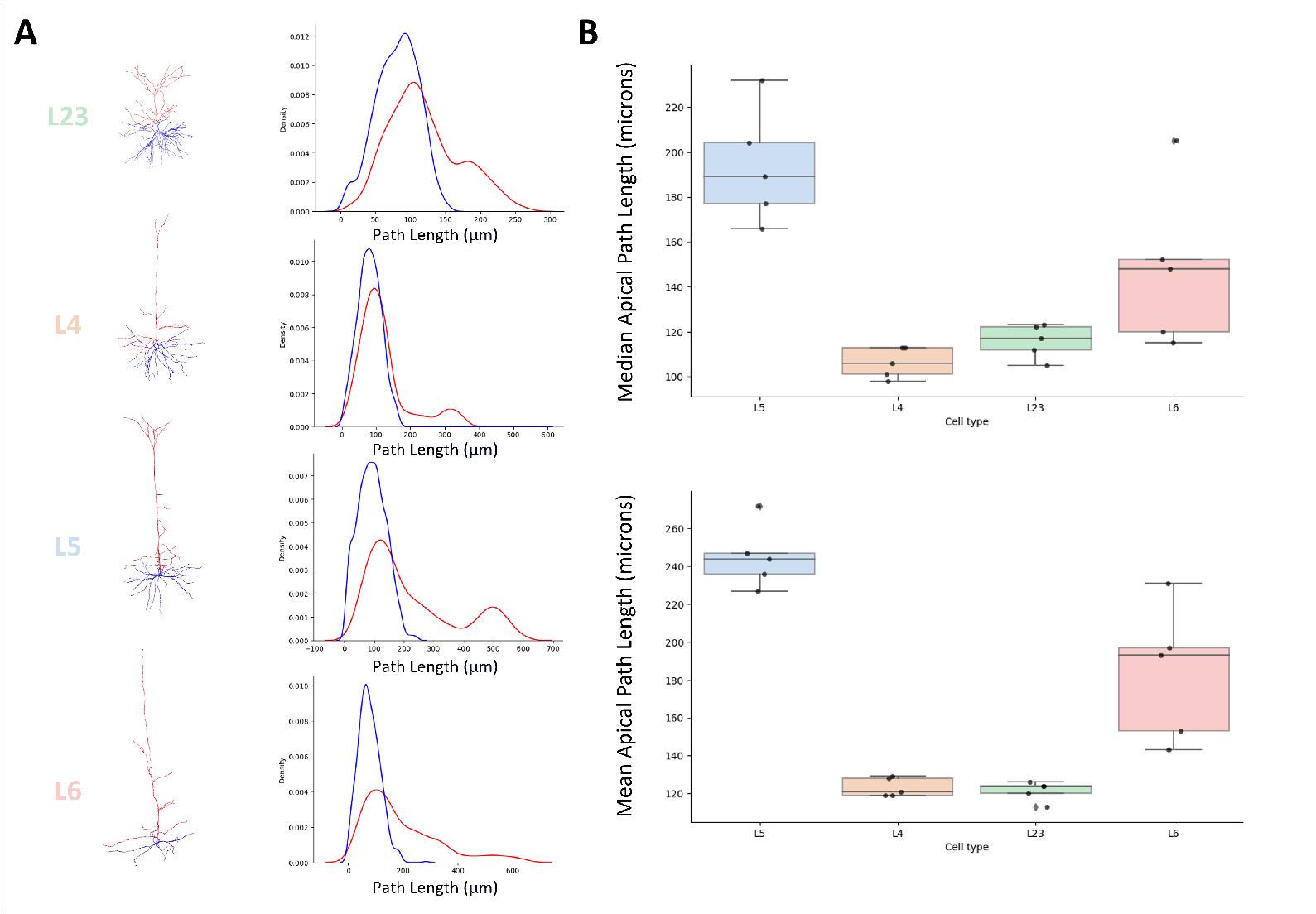
Apical synapses across neuron types are predominantly local, with layer-specific variation in path length. (A) Example reconstructions (left) and corresponding density plots (right) showing synaptic path lengths from apical synapses back to the soma for neurons from layers L2/3, L4, L5, and L6. Blue curves represent basal synapses for comparison; red curves show apical synapse path lengths. (B) Quantification of apical path lengths across 5 neurons per class. Top: Median apical path length; Bottom: Mean apical path length. Layer 5 neurons exhibit the longest apical synaptic paths, while L2/3 and L4 neurons show more compact, local apical arborizations. Variability in L6 reflects heterogeneity in apical morphology.

**Figure S3:**
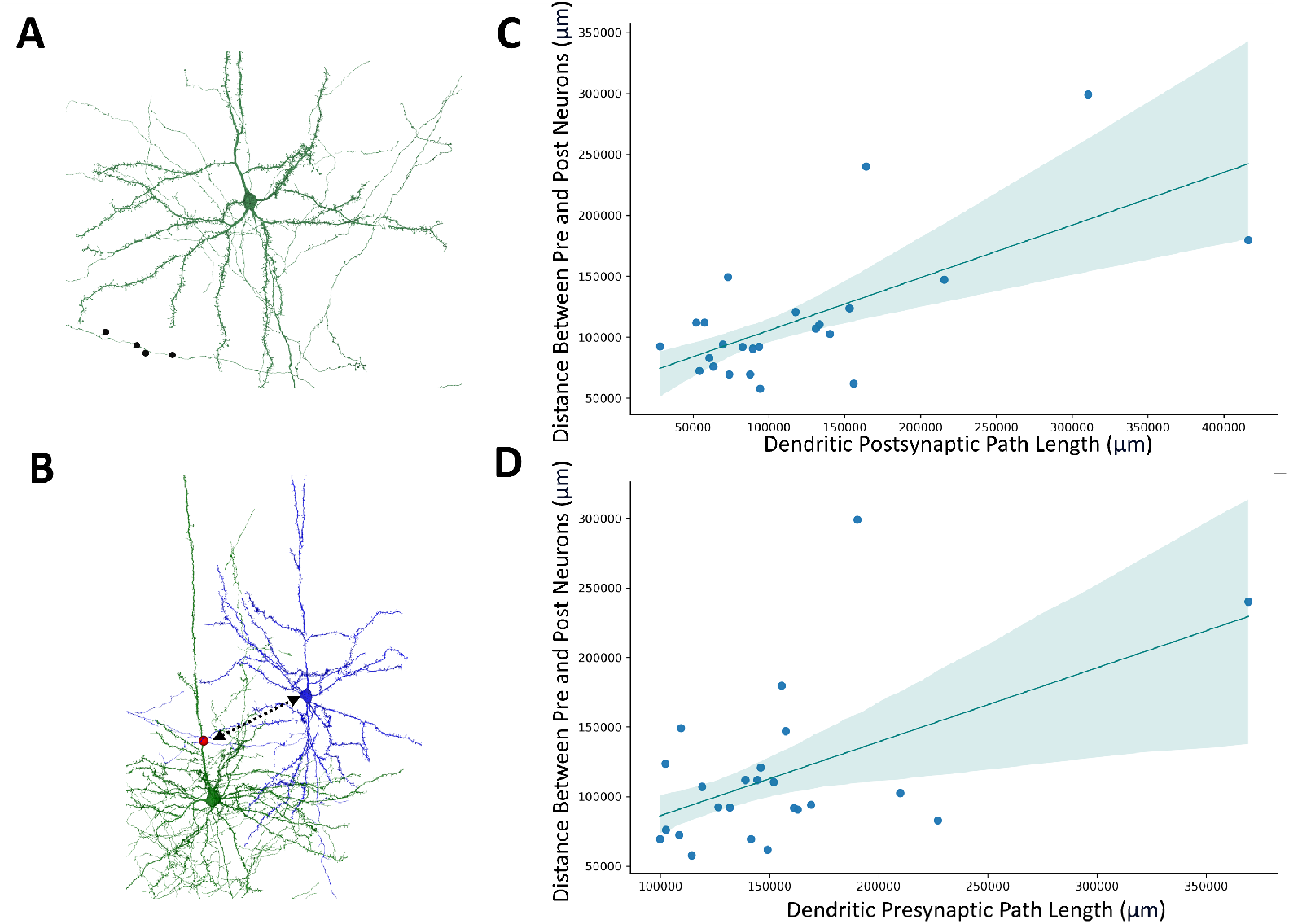
Path length from soma predicts intersomatic distance for both pre- and postsynaptic neurons. (A) Example manually traced axon (green) with synapses (black dots) used to identify postsynaptic targets. (B) Example of a synaptically connected neuron pair: presynaptic (green) and postsynaptic (blue) neuron morphologies are shown, with the synapse indicated (red). (C) Euclidean distance between connected neuron somas is significantly correlated with dendritic path length from the synapse to the postsynaptic soma (*p <* 0.0001). (D) A similar relationship holds for path length from the presynaptic soma to the synapse (*p* = 0.0001). Shaded regions represent 95% confidence intervals from linear regression.

**Figure S4:**
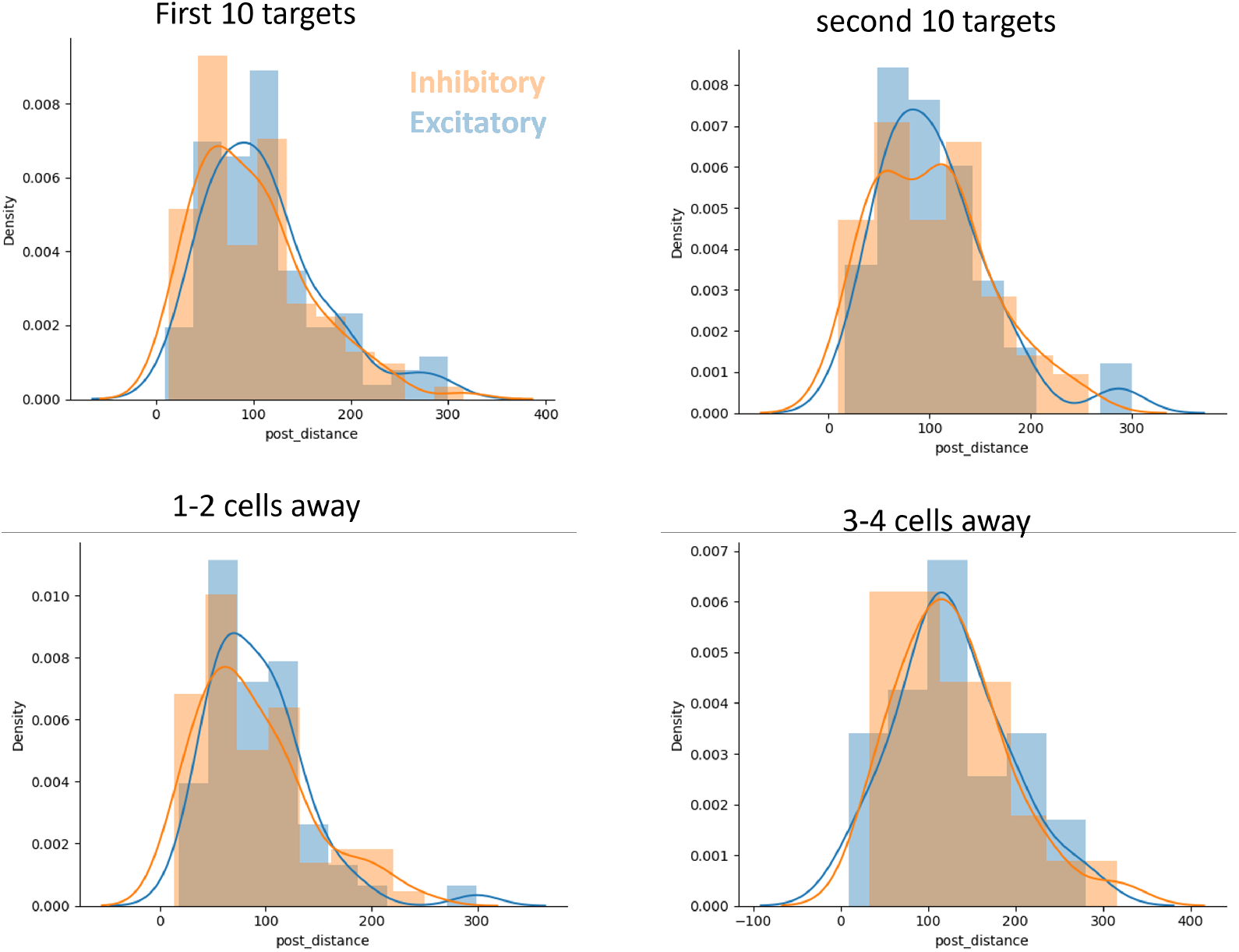
Postsynaptic dendritic path lengths do not significantly differ between excitatory and inhibitory target neurons. Density histograms showing the distribution of dendritic path lengths from synapse to soma (in microns) for postsynaptic targets categorized as excitatory (blue) or inhibitory (orange). Each subplot represents a different subset of the data. Across all comparisons, there were no statistically significant differences in path length distributions between excitatory and inhibitory postsynaptic neurons.

**Figure S5:**
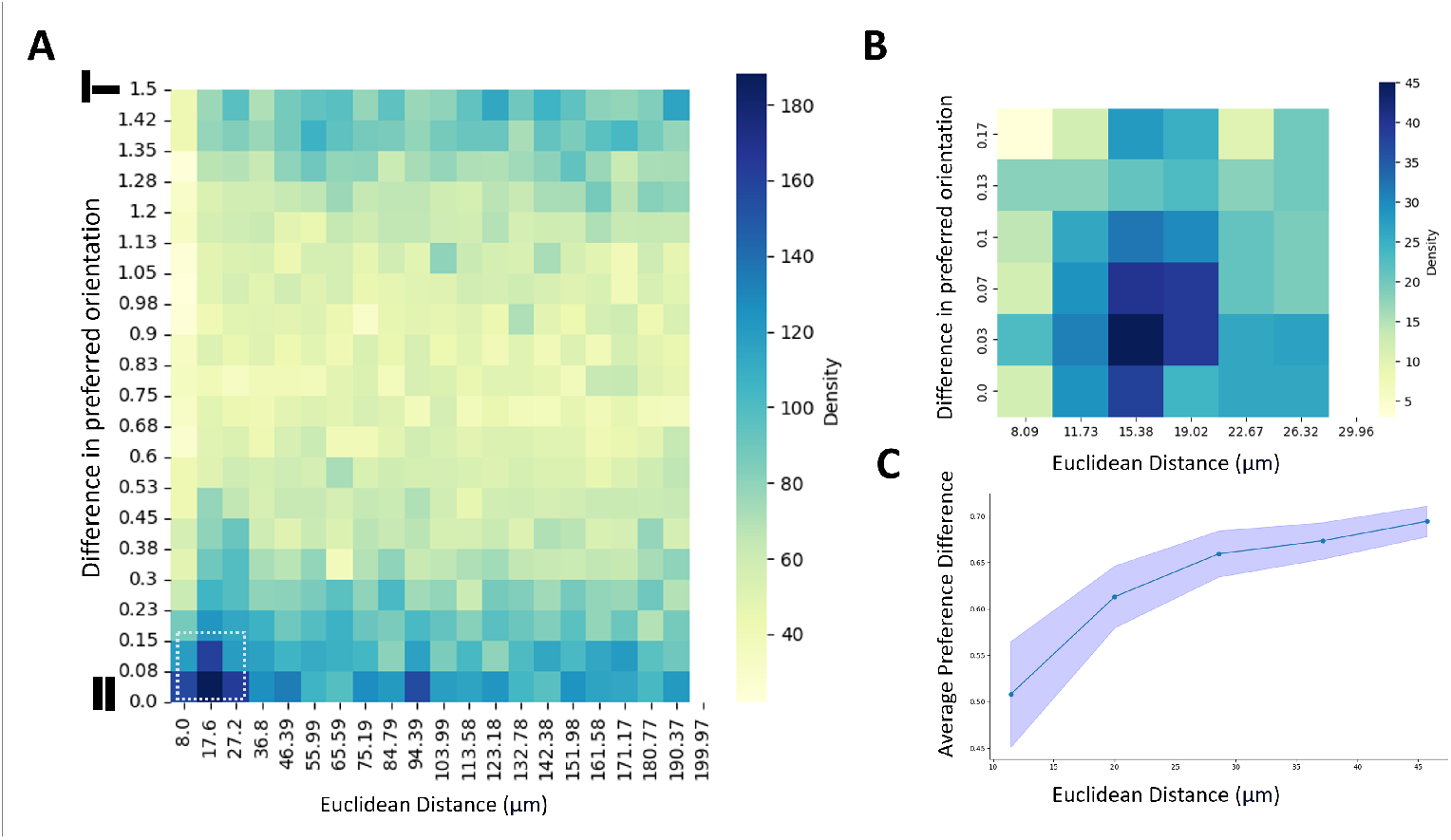
Nearby neurons exhibit similar stimulus preferences, consistent with proximity-based connectivity. (A) Joint density plot showing the relationship between Euclidean distance and difference in preferred orientation for neuron pairs in the MiCRONS dataset. Neuron pairs that are closer in space tend to exhibit more similar stimulus tuning. (B) Magnified view of the region highlighted in (A), showing higher density of neuron pairs with both close spatial proximity and small differences in preferred orientation. (C) Average difference in preferred orientation as a function of Euclidean distance. Neurons located within *∼*20 *µ*m show significantly more similar stimulus tuning than those farther apart. Shaded region indicates SEM. These results support the hypothesis that structural proximity enhances functional similarity among connected neurons.

## Notes

### Competing Interest Statement

The authors have declared no competing interest.

